# Hydroquinone, a cigarette smoke compound, affects cartilage homeostasis through activation of the aryl hydrocarbon receptor pathway

**DOI:** 10.1101/2022.04.25.489372

**Authors:** Cintia Scucuglia Heluany, Anna de Palma, Nicholas Day, Sandra Helena Poliselli Farsky, Giovana Nalesso

## Abstract

Exposure to cigarette smoke has a proven detrimental impact on different aspects of human health. Increasing evidences link smoking to degeneration of joint tissues. However, the toxic mechanisms elicited by the different components of cigarette smoke have not been fully elucidated yet. We have previously shown that exposure to hydroquinone (HQ), a pro-oxidant chemical present in cigarette smoke, can promote joint tissue degradation in murine models of rheumatoid arthritis through the activation of the aryl hydrocarbon receptor (AhR) pathway. Osteoarthritis (OA) is a chronic debilitating articular disease characterized by progressive degradation of the articular cartilage, whose onset and progression have also been associated with smoking. In this work we aimed to investigate the effect of HQ exposure on articular chondrocytes and how it affects cartilage homeostasis. Cell viability, gene expression, oxidative stress and inflammatory parameters were quantified in primary articular chondrocytes exposed to HQ in presence or absence of IL-1β pre-stimulation. HQ stimulation downregulated phenotypic markers genes such as SOX-9 and Col2a1, whereas upregulated the expression of the catabolic enzymes MMP-3 and ADAMTS5. HQ also promoted oxidative stress and reduced proteoglycan content. HQ exacerbated the pro-inflammatory effects mediated by the IL-1β co-stimulation. Finally, we showed that HQ-degenerative effects were mediated by the activation of AhR. Together, our findings address the harmful effects of HQ in the articular cartilage health, providing novel evidence surrounding the toxic mechanisms of environmental pollutants underlying the onset of articular diseases.

## Introduction

The exposure to environmental pollutants can severely affect and compromise human health. Cigarette smoke is considered the major environmental risk factor for the developmental and progression of chronic inflammatory diseases such as chronic obstructive pulmonary disease (COPD), cardiovascular diseases and psoriasis (Stampfli and Anderson, 2009).

Hydroquinone (HQ) is a chemical compound representing 3 % of the particle matter phase of cigarette smoke (Bodnar et al., 2012). This xenobiotic is a benzene metabolite which can mediate immune- and mieolotoxicity (Mcgregor, 2007; Li et al., 2018). Exposure to HQ has been also associated with increased apoptosis and development of oxidative stress in immune cells (Cho et al., 2008; Lee et al., 2012). In vitro studies showed that HQ alone is responsible for 10 % of oxidative stress-mediated cellular toxicity when cells are exposed to cigarette smoke (Stabbert et al., 2017).

We have recently shown that exposure to HQ aggravated joint damage in experimental animal models of rheumatoid arthritis (RA) through the activation of the aryl hydrocarbon receptor (AhR) pathway by promoting inflammation, an increased influx of immune cells, synovial proliferation and oxidative stress (Heluany et al., 2018a,b; Heluany et al., 2021).

Exposure to cigarette smoke has also been associated with increased pain and cartilage loss in both osteoarthritis (OA) and in degenerative disc diseases (Goldeberg, Scott and Mayo, 2000; Fogelholm and Alho, 2001; Amin et al., 2007). So far, the main research focus has been the investigation of the role of nicotine in the disease progression, often with inconclusive or contrasting results (Felson and Zhang, 2015).

OA is the most widespread and disabling type of chronic and degenerative articular disease, characterized by low-grade inflammation, breakdown of the articular cartilage, abnormal remodeling of the subchondral bone and growth of bone spurs, also known as osteophytes (Berenbaum, 2013; Mobasheri et al., 2017). The chondrocyte is the only cell type populating the articular cartilage and plays an essential role in maintaining cartilage homeostasis. OA progression is tightly linked to the loss of chondrocyte function, due to the deregulation of several signaling cascades and the acquisition of a hypertrophic phenotype (Akkiraju and Nohe, 2015; Thomas et al., 2007). In addition, the upregulation of pro-inflammatory cytokines such as interleukin-1β (IL-1β) has been strongly associated with loss of chondrocyte homeostasis and exacerbation of articular cartilage damage in OA (Sellan and Berenbaum, 2010).

In this work we investigated the effect of HQ on articular chondrocytes to elucidate how it can affect the articular cartilage health. We stimulated bovine articular chondrocytes and cartilage explants with HQ to investigate its effects on cell viability, modulation of phenotypic genes and oxidative stress. Our results showed that HQ exposure altered cell proliferation, down-regulated the expression of the phenotypic markers SRY-Box transcription factor 9 (SOX-9), type II collagen (Col2a1) and type X collagen (Col-X), while up-regulating the expression of catabolic markers matrix metalloprotease 3 (MMP-3) and A Disintegrin and Metalloproteinase with Trombospondin motifs 5 (ADAMTS5). HQ exposure also promoted oxidative stress and potentiated the pro-catabolic effects of IL-1β. We also showed that HQ activity is mediated through the activation of the AhR signaling pathway. Together, our findings suggest that the AhR activation in articular chondrocytes could be an important mechanism through which environmental pollutants compromise cartilage homeostasis.

## Material and Methods

### Isolation and culture of bovine articular chondrocytes and cartilage explants

Bovine articular cartilage explants were isolated from the metatarsal and metacarpal joints of adult cows purchased within less than 24 hours of death from a local abattoir. The articular cartilage was dissected in sterile conditions and processed for chondrocyte isolation or explant cultures as previously described (Nalesso et al., 2011). Bovine articular chondrocytes (BACs) were cultured with complete medium (CM; DMEM F12 supplemented with 10 % fetal bovine serum (FBS) and 1 % of antibiotics penicillin and streptomycin) at 37 °C in a 5 % CO_2_ incubator. Cells at passage 1 to 3 were used for the experiments described in this manuscript.

### MTT assay

Cell integrity and viability were measured through the 3-(4,5-dimethylthylthiazol-2-yl)- 2,5 diphenyltetrazolium bromide (MTT) method. Briefly, 1 × 10^4^/well BACs were seeded in 96-well plates and cultured in CM for 24 hours. Thereafter, cells were washed with PBS and cultured with DMEM F12 supplemented with 0.1 % of BSA for 24 hours. BACs were treated as described in the individual experiments. Then, 0.5 mg/mL of MTT solution was added to each well and incubated for 3 hours in the dark at 37 °C. Thus, the medium was removed, and the blue formazan crystals were dissolved in 200 µL of DMSO. The optical density reading was recorded at 570 nm in a plate reader (Clario Star, BMG LABTECH).

### Gene expression

For gene expression analyses, 7.5 × 10^4^ BACs were plated per well in 24-well plates and cultured with CM for 24 hours. Then the cells were washed with PBS and treated as described in the Results section. Total RNA was extracted from BACs using TRIzol reagent (Invitrogen), according to the manufacturer’s instructions. Three hundred and fifty nanograms of total RNA from each sample were reverse transcribed to cDNA using a High Capacity cDNA Reverse Transcription Kit (Applied Biosystems). Quantitative PCR was performed with hot-start DNA polymerase (Qiagen), as previously described (Nalesso et al., 2011). Primer sequences are listed in Table 1. All data were normalized to internal control of β-actin values. All experiments were performed in a BioRad PCR system (CFX96TM Optics Module).

**Table 1:**
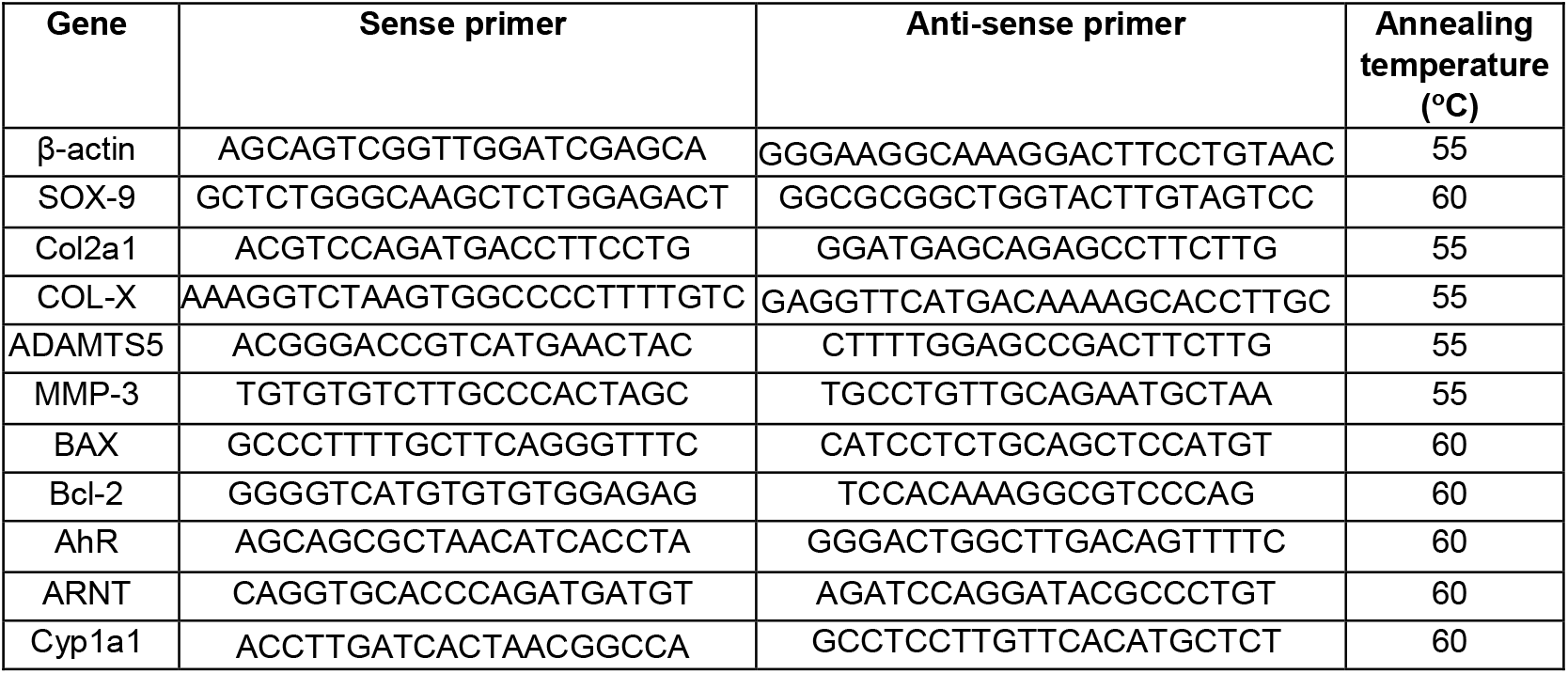
PCR primer sequences used for quantitative gene expression analysis.

### Micromass cultures

BACs were plated at a density of 25 × 10^4^ cells per well in complete medium in 24-well plates. Micromass cultures were prepared as described before (De Bari et al., 2001). The micromasses were stimulated as described in the individual experiments with HQ ± IL-1β (20 ng/mL; RP0106B-025 Kingfisher Biotech). After 48 hours, the culture media were collected and the micromasses were washed with PBS and fixed with cold methanol.

### Alcian blue staining

The micromasses were stained overnight with Alcian blue, as previously described (De Bari et al., 2001). After proteoglycan extraction with 6 M guanidine hydrochloride (Sigma-Aldrich), the absorbance was read at 630 nm with a spectrophotometer (Clario Star, BMG LABTECH). The absorbance values were normalized to protein content. Images of the micromasses were acqurided at room temperature with a stereomicroscope (SZTL 350 Stereo Binocular Microscope, VWR®).

### Safranin-O staining

Bovine articular cartilage explants were stimulated ex vivo with HQ as described in the Results section. At the end of the exposure period after 72 hours, the culture media was collected while the cartilage specimens were fixed in 4 % paraformaldehyde (PFA) and paraffin embedded. Three-micrometre thick sagittal sections were stained with safranin-O as described before (Nalesso et al., 2011), and images were taken using the same settings on an optical microscope (Nikon). Images were acquired through 10x magnification objective lenses and the SO-staining was quantified by using the ImageJ software (NIH). For such, 4 fields per slide (totalling 16 images per experimental group) were considered for the analysis and the mean of intensity from all groups was generated and used for the statistical analysis.

### DMMB assay

Release of glycosaminoglycans (GAGs) by micromasses and cartilage explants upon exposure to HQ was measured in the culture media by using the Dimethyl methylene Blue (DMMB) assay (adapted from Farndale et al., 1982). Briefly, 200 µL of the DMMB solution was added to 20 µL of the culture media in a 96-well plate. Subsequently, the absorbance was measured at 525 nm with a spectrophotometer (Clario Star, BMG LABTECH).

### Quantification of nitric oxide by Griess assay

Fifty microliters of Sulphanilamide Solution (1 % in 5 % phosphoric acid; Promega) was added to 50 µL of the culture media from micromass cultures of BACs or bovine cartilage explants treated with HQ ± IL-1β (20 ng/mL). After a 10 minutes incubation at RT, 50 µL of N-1-napthylenediamine dihydrochloride solution (NED, 0.1 % in water; Promega) was added to the mix. After a further 10 minutes incubation, the absorbance was measured at 550 nm with a spectrophotometer (Clario Star, BMG LABTECH).

### Quantification of Reactive oxygen species by DCFH

The intracellular accumulation of reactive oxygen species (ROS) was quantified in BACs cultures using the fluorescent probe CM-H2DCFDA. Ten thousand chondrocytes/well were seeded in 24-well plates and stimulated with HQ for 24 hours. The cells were then incubated with 10 μM CM-H2DCFDA for 30 minutes at 37 °C in the dark. The cells were resuspended in PBS and 10,000 events were acquired in a MACSQuant flow cytometer. Results are presented as arbitrary units of fluorescence.

### Reporter assay

BACs were seeded at a density of 7 × 10^4^ cells/well in CM on a 24-well plate. The cells were co-transfected with 450 ng/well of pGL4.43 [luc2P/XRE/Hygro] luciferase reporter vector (Promega) and with 50 ng/well of the control vector expressing *Renilla reniformis* luciferase by using Lipofectamine (Thermo Fisher) following the manufacturer instructions. After the transfection, the cells were stimulated with CM (vehicle), or with the AhR ligand 6-formylindolo(3,2b)carbazole (FICZ, 10 µM) or with HQ (10 or 25 µM) for 24 hours. Luciferase activity was determined using the Dual luciferase reporter assay system (Promega). Firefly luciferase activity was normalized by the Renilla luciferase activity. Data are expressed as fold increase of relative luminescence units in comparison to vehicle.

### Statistical analysis

One-way ANOVA with Tukey-Post test was used to compare the statistical differences between multiple groups, and a two-tailed *t*-test was used for comparisons between two groups. Values of p < 0.05 were considered statistically significant. Data are expressed as the mean ± standard error of the mean (SEM). Statistical analyses were performed using GraphPad Prism version 8.0 (GraphPad Software, CA, USA).

## Results

### In vitro exposure to HQ halts articular chondrocyte growth but does not induce apoptosis

To investigate whether exposure to HQ could affect chondrocyte viability, we treated bovine articular chondrocytes with different concentrations of HQ (1 μM up to 100 μM) and monitored cell growth across time by MTT assay.

HQ treatment slowed cell growth at concentrations between 10 μM to 100 μM over a 96h time course in comparison to vehicle control (Figure 1A). While affecting cell growth, HQ did not alter cell viability under the concentrations of 1 to 50 µM observed at the 72h time point (Figure 1B) and did not induce activation of apoptosis, as shown by the unaltered ratio of Bax/Bcl2 expression at mRNA levels (Figure 1C) (Karaliotas et al., 2015).

**Figure 1:**
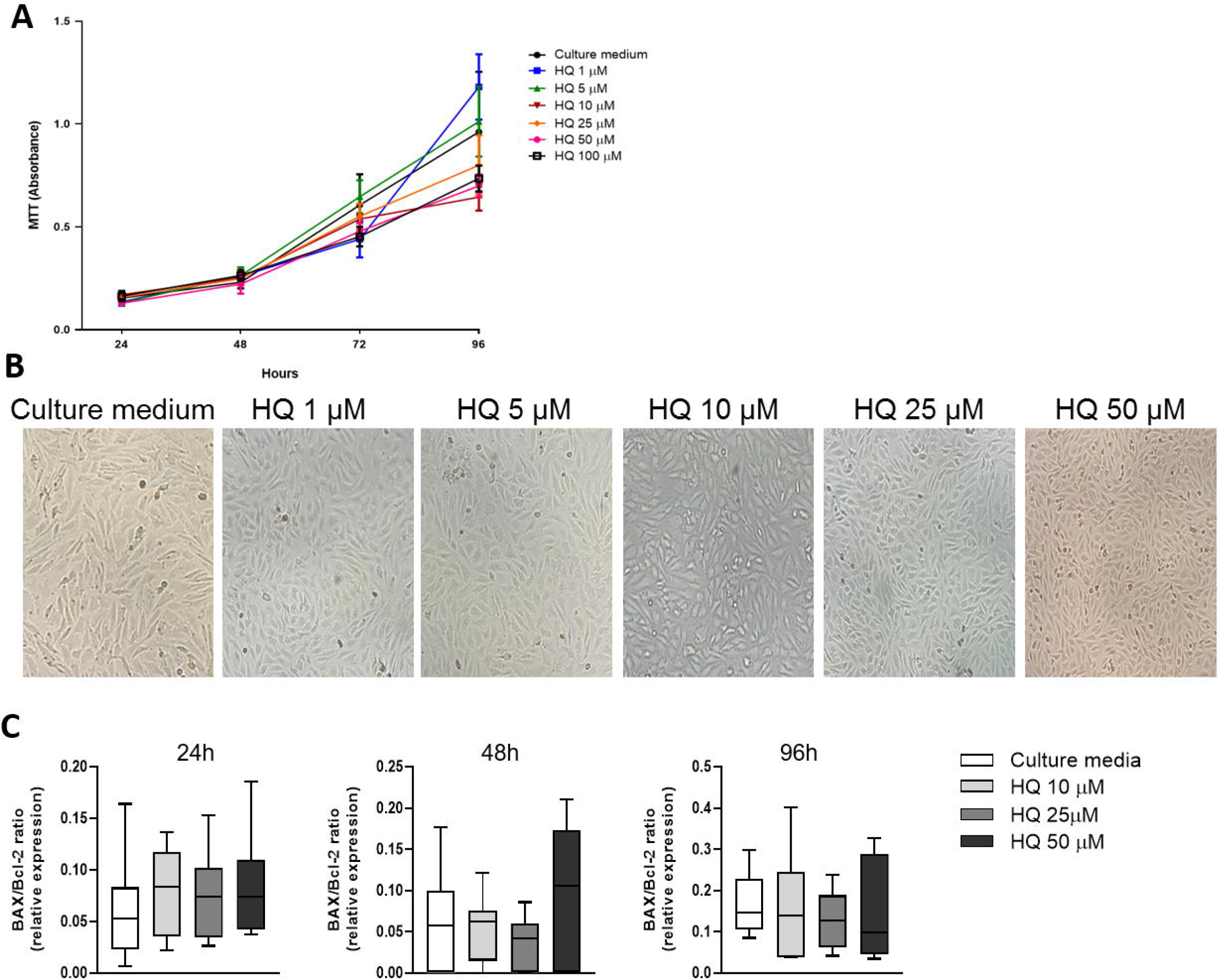
In vitro HQ exposure reduces viability but does not promote cell death. Stimulation of BACs with increasing concentrations of HQ (1-100 µM) halted cell growth as evaluated by MTT assay **(A**) and presented in representative images after 72 hours of treatment **(B)**. HQ treatment did not promote apoptosis **(C)** as the BAX/Bcl-2 mRNA ratio was not different across treatments and over time. Data represent mean ± SEM from three independent experiments and were analysed by one-way ANOVA. 16H: p= 0.0037 CM *vs*. 100 µM. 24H: p= 0.0215 CM *vs*. 1 µM; p= 0.0220 CM *vs*. 5 µM; p= 0.0010 CM *vs*. 10 µM; p= 0.014 CM *vs*. 25 µM; p = 0.0001 CM *vs*. 50 and 100 µM. 72H: p= 0.0049 CM *vs*. 25 µM; p = 0.0001 CM *vs*. 50 and 100 µM. 96H: p= 0.0262 CM *vs*. 10 µM; p = 0.0001 CM *vs*. 25, 50 and 100 µM.

### HQ modulates the expression of phenotypic markers in articular chondrocytes

We further investigated whether HQ treatment could influence the expression of phenotypic markers in articular chondrocytes, as these markers are altered during the progression of OA (Dell’Accio et al., 2001; Nalesso et al., 2011; Nalesso et al., 2017).

The mRNA expression of the differentiation markers SOX-9, Col2a1 and Col-X was significantly downregulated upon incubation with HQ for 24 hours (Figure 2A-C). Furthermore, the expression of the matrix remodelling enzyme MMP-3 and of the aggrecanase ADAMTS5 was instead upregulated (Figure 2D, E). The catabolic activity of HQ was further confirmed by a reduced amount of highly sulphated GAGs in chondrocyte micromasses upon treatment with HQ (Figure 2F, G).

**Figure 2:**
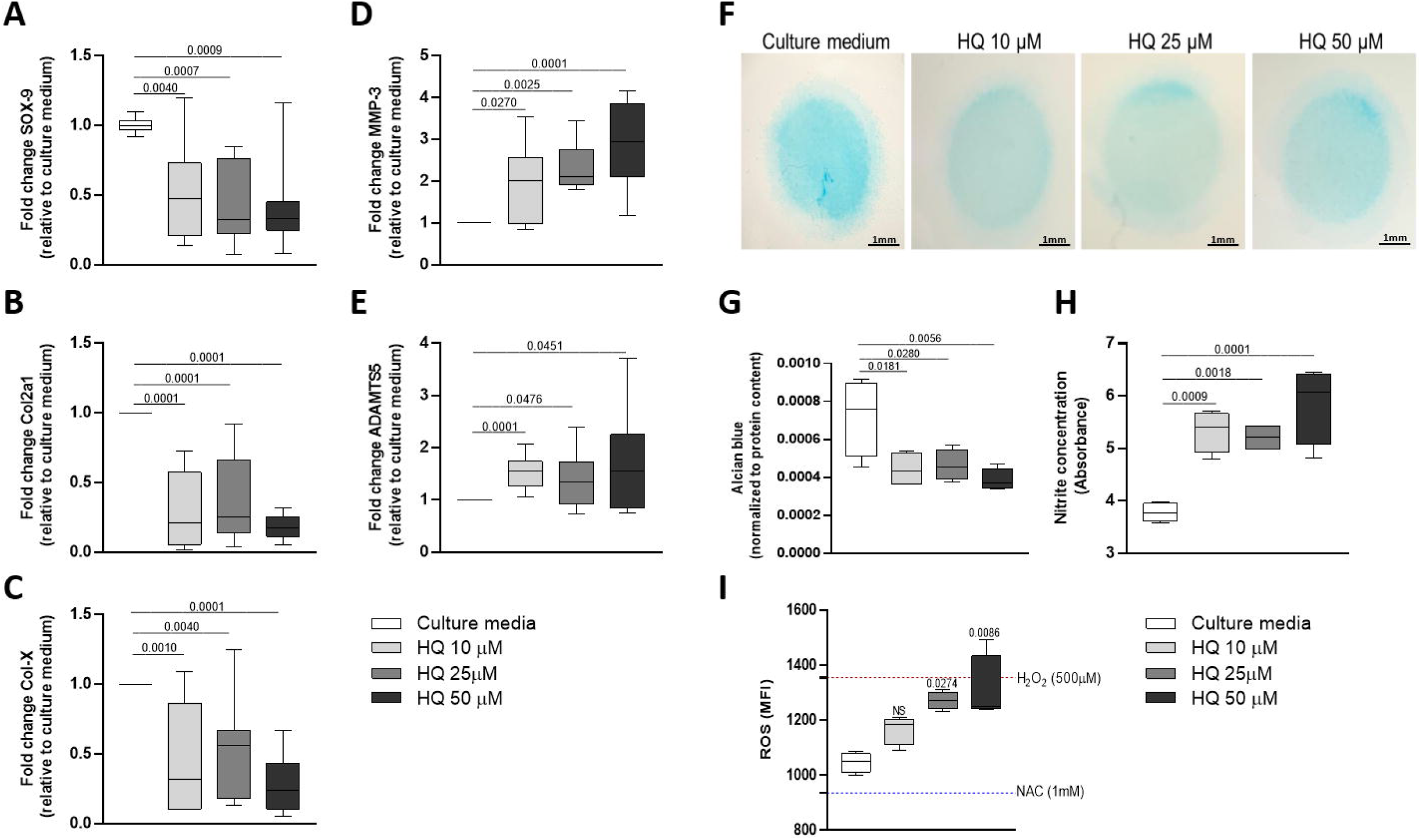
HQ exposure modulated phenotypic markers in articular chondrocytes. HQ stimulation of monolayer cultures of BACs downregulated the phenotypic markers SOX-9 **(A)**, Col2a1 **(B)**, Col-X **(C)** but upregulated the catabolic enzymes MMP-3 **(D)** and ADAMTS5 **(E)**. HQ exposure reduced the production of highly sulphated GAGs **(F-G)** over a 48h incubation on BACs in micromass cultures, as measured by Alcian Blue quantification. HQ also increased nitrite production (**H**, quantified by Griess reaction) and ROS generation (**I**, quantified by DCFH-DA assay) in micromass cultures. qPCR results were normalized to the housekeeping gene β-actin and expressed as fold change to culture media. Data represent mean ± SEM from three independent experiments and were analysed with one-way ANOVA and unpaired t test.

Nitric oxide (NO) can suppress the synthesis of proteoglycans (PGs) and stimulates MMPs activity (Amin and Abramson, 1998). We therefore tested whether HQ could elicit NO production in articular chondrocytes. Indeed, HQ exposure for 48 hours significantly increased the nitrite production, as evaluated by Griess reaction (Figure 2H).

Finally, as HQ can induce the generation of ROS in different biological systems (Heluany et al., 2018b; Heluany et al., 2021; Mao et al., 2019), and the induction of oxidative stress in chondrocytes has been strongly associated with cartilage damage in OA (Henrotin, Bruckner and Pujol, 2003), we tested whether the HQ exposure could influence the production of ROS in BACs cultures. Our data showed that an incubation of chondrocytes with HQ for 24 hours triggered a significant increase in the levels of ROS in comparison to non-stimulated cells (Figure 2I).

Altogether, these results suggest that HQ alone can promote alteration in the chondrocyte metabolism and oxidative stress, which are associated with activation of tissue degenerative mechanisms.

### HQ enhanced the pro-catabolic activity of IL-1β

IL-1β is a pro-inflammatory cytokine overexpressed in the articular cartilage during the progression of OA, promoting cartilage degradation processes (Sellan and Berenbaum, 2010). Thus, to investigate whether inflammation could influence the catabolic activity of HQ, BACs were pre-stimulated with IL-1β at 20 ng/mL for 4 hours before being exposed to different concentrations of HQ for 24 hours or 48 hours. While HQ does not exacerbate the pro-catabolic effects of IL-1β at gene expression levels (Figure 3A, B), pre-stimulation with IL-1β increased GAG content reduction and NO generation induced by HQ, in a dose dependent fashion (Figure 3C-F).

**Figure 3:**
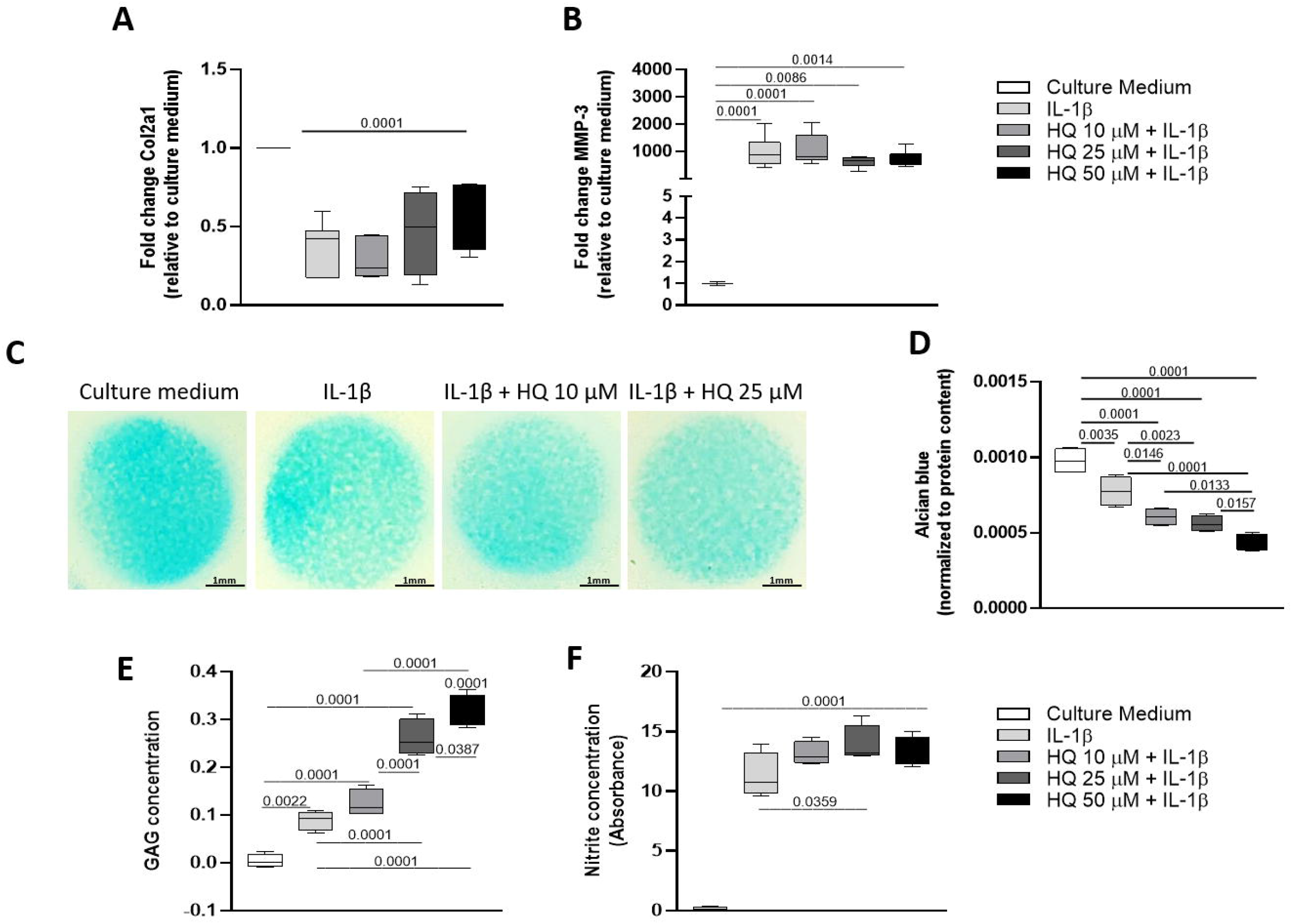
HQ treatment enhances the pro-catabolic activity of IL-1β in articular chondrocytes. Pre-incubation of monolayer cultures of BACS with IL-1β (20 ng/mL) for 4 hours before stimulation with HQ (10, 25 or 50 µM) for 48 hours, did not alter the expression of the phenotypic marker Col2a1 **(A)** and of the catabolic enzyme MMP-3 **(B)** at mRNA level. Co-stimulation of IL-1β and HQ reduced the accumulation of highly sulphated GAGs **(C**-**D)**, as assessed by quantification of Alcian blue staining. The co-treatment also enhanced the release of GAGs **(F)** and increased the nitrite production **(G)** in micromass cultures of BACs, as measured respectively by DMMB assay and Griess reaction, respectively. qPCR results were normalized to the housekeeping gene β-actin and expressed as fold change to culture media. Data represent mean ± SEM from three independent experiments and were analysed with one-way ANOVA and unpaired t test.

These results were further confirmed on ex-vivo cultures of bovine cartilage explants. While the IL-1β pre-treatment did not influence the GAG content and the release induced by HQ (Figure 4A-B, D), HQ and IL-1β showed to have a cumulative effect on nitrite production (Figure 4C).

**Figure 4:**
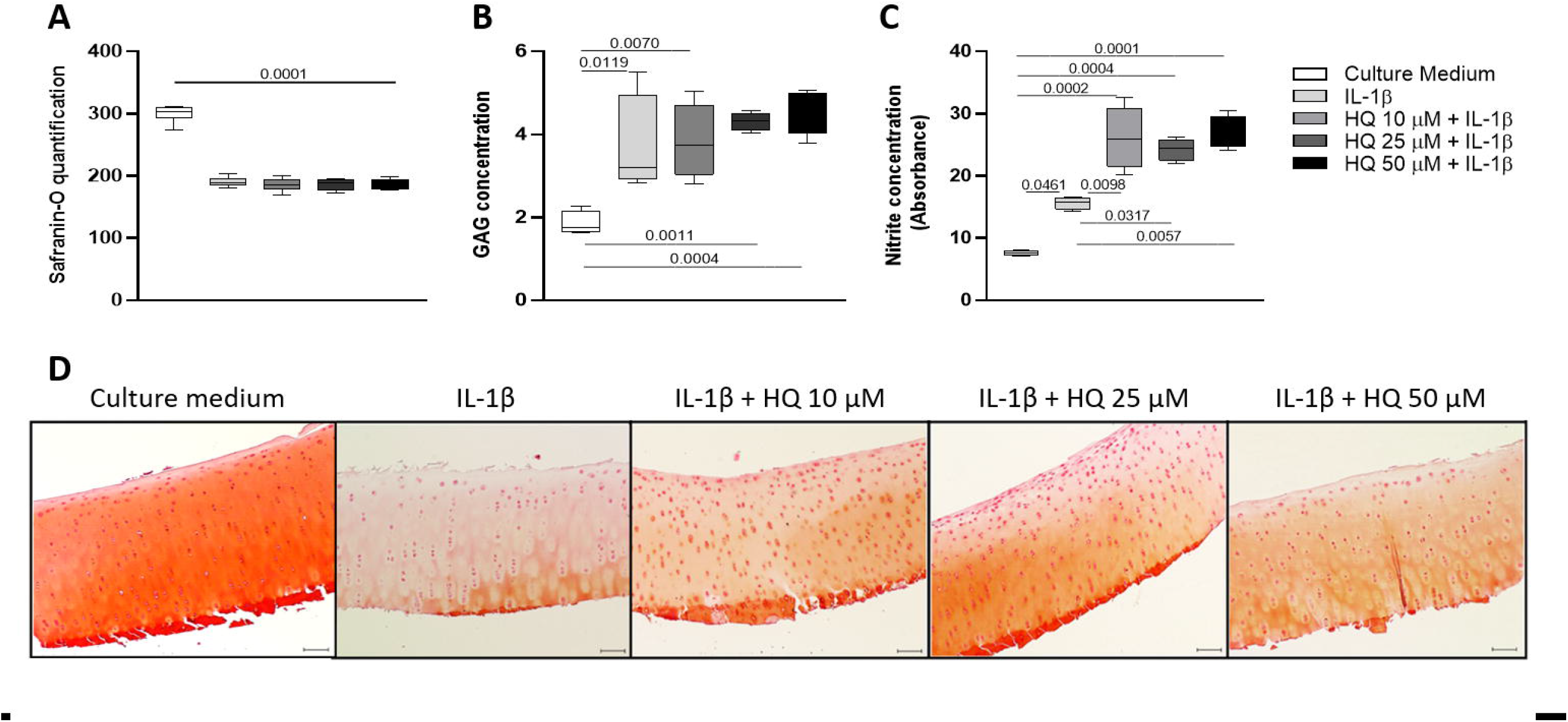
HQ treatment increases the pro-inflammatory activity of IL-1β in articular cartilage explants. Bovine articular cartilage explants were pre-incubated with CM or with IL-1β (20 ng/mL) for 4 hours and subsequently incubated with HQ (10, 25 or 50 µM) for 72 hours. The co-stimulation did not alter GAGs synthesis and release **(A-B, D)**, but increased nitrite production **(C)** in comparison to stimulation with IL-1β alone. 10X magnification. Data represent ± SEM from four independent experiments and were analysed with one-way ANOVA.

Altogether, our findings suggest that the pro-inflammatory stimuli can exacerbate the catabolic effects of HQ in the articular cartilage.

### HQ activates the AhR pathway in articular chondrocytes

We previously demonstrated that HQ mediates its toxic effect in the joints through activation of the AhR pathway in murine experimental models of rheumatoid arthritis (Heluany et al., 2018a,b; Heluany et al., 2021). Thus, to investigate whether HQ signals through this pathway in the articular chondrocytes, we measured the expression levels of AhR, of the aryl hydrocarbon receptor nuclear translocator (ARNT) and of the endpoint target gene in the AhR pathway Cyp1a1 (Figure 5A) in BACS upon exposure to different concentrations of HQ. All the genes were upregulated in chondrocytes upon incubation with HQ for 24 hours and their modulations were rescued by the pre-incubation with α-naphthoflavone (αNF, 10 µM), an AhR antagonist (Figure 5B-D). (Nguyen et al., 2015).

**Figure 5:**
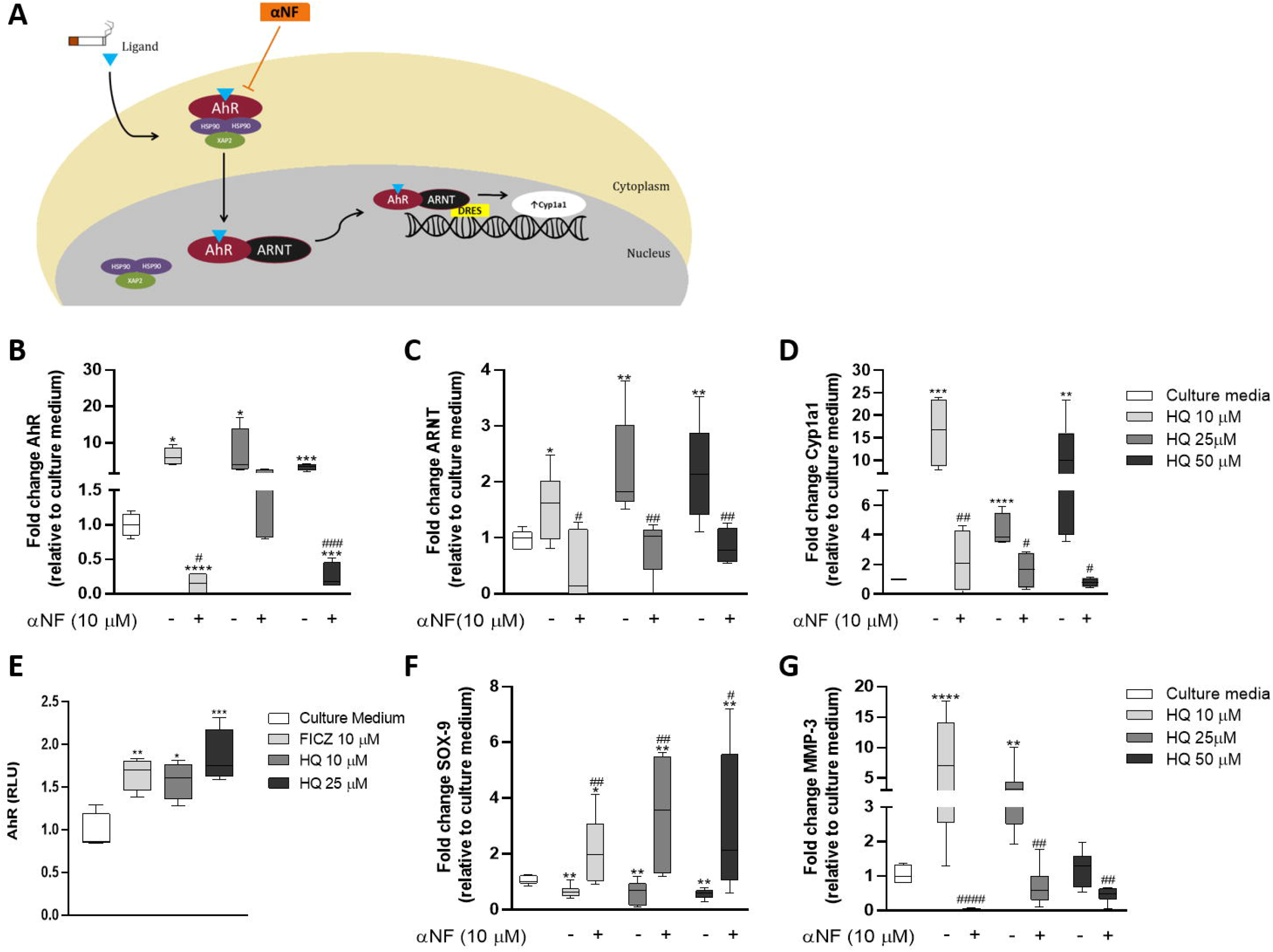
HQ activates the AhR pathway in articular chondrocytes: **(A)** Schematic representation of the AhR pathway. Modulation of components of the AhR pathway was determined by measuring mRNA levels of AhR, ARNT and Cyp1a1 in BACs incubated with CM or with HQ (10, 25 or 50 µM) in absence or presence of αNF (10 µM) for 24 hours **(B-D)**. Results were normalized to the housekeeping gene β-actin and are expressed as fold change to CM. **(E)** Activation of the AhR receptor by HQ was confirmed through a luciferase-based reporter assay. Inhibition of AhR with αNF rescued the downregulation of SOX-9 **(F)** and the upregulation of MMP-3 **(G)** mediated by HQ. Data represent mean ± SEM from three independent experiments and were analysed by one-way ANOVA. *p<0.05, **p<0.01, ***p<0.001 and ****p<0.0001 *vs*. CM; ^#^p<0.05, ##p<0.01, ###p<0.001 and ####p<0.0001 *vs*. IL-1β.

In the activated stage, the AhR translocates from the cytoplasm to the nucleus and forms a complex with the ARNT, which identifies and integrates with the xenobiotic responsive elements (XRE) of target genes, regulating their expression (Ma and Baldwin, 2000). Activation of the pathway upon HQ stimulation was therefore further confirmed with an XRE-reporter assay in articular chondrocytes stimulated for 24 hours with HQ. As shown in the Figure 5E, treatment with HQ increased the intensity of the binding activity to the AhR gene reporter in a concentration-dependent manner. Finally, we confirmed that both the phenotypic changes as well as the pro-catabolic activity induced by HQ were mediated via the activation of the AhR pathway, as the HQ-mediated downregulaton of SOX-9 and upregulation of MMP-3 were rescued by the pre-incubation with the AhR antagonist αNF (Figure 5F, G).

## Discussion

The effect of cigarette smoke on the health of joint tissues has not been fully elucidated yet. In a few observational studies, exposure to cigarette smoke has been shown to trigger contrasting effects, as smoking can both increase or reduce the incidence of OA, depending on the cohort examined and the methodology used for the analysis (Davies-Tuck et al., 2009; Racunica et al., 2007).

In an attempt to further elucidate how cigarette smoke can impact the health of joint tissues at molecular levels, here we focused our attention on investigating the role of HQ, a major component of cigarette smoke on articular cartilage health, as we have previously shown its contribution to the degeneration of other tissues in the joints in murine models of inflammatory arthritis (Heluany et al., 2018a,b; Heluany et al., 2021).

Our data showed that HQ decreases chondrocyte viability in a dose and time dependent manner. We and others previously associated the toxic effect of HQ with activation of apoptosis in different biological systems (Lee et al., 2012; Li et al., 2018), however, we did not detect any variation in apoptotic markers in articular chondrocytes upon HQ exposure within the timeframe of our analysis.

In OA cartilage, the articular chondrocytes undergo phenotypic changes, characterized by the downregulation of phenotypic lineage markers and the upregulation of pro-hypertrophic genes, as well as of matrix remodelling enzymes, which contribute to the progressive loss of the biomechanical properties of the tissue, and ultimately lead to its degradation (Arner, 2002; Poole et al., 2001). Our data suggest that HQ could heavily contribute to many of these phenomena. We showed that HQ exposure promoted the downregulation of phenotypic markers, while also inducing the upregulation of MMP-3, a metalloprotease, whose activity is upregulated in OA, which can degrade several types of collagens and matrix proteins in the articular cartilage (Bortoluzzi et al., 2018). The enhancement of matrix remodelling activity induced by HQ was also supported by the reduction of GAG staining and increased GAG release upon exposure of chondrocytes 2D cultures to the xenobiotic.

Oxidative stress can be induced through several mechanisms and can contribute to cellular metabolic decline as well as to promote degenerative mechanisms (Suantawee et al., 2013). The induction of oxidative stress in chondrocytes has been strongly associated with increased cartilage degradation in severe cases of OA (Henrotin, Bruckner and Pujol, 2003). Moreover, nitrite production has also been shown to contribute to oxidative stress and degeneration of tissue integrity in the joints (Abramson, 2008). Previous data showed that exposure to environmental pollutants could promote ROS and NO generation in chondrocytes (Lee and Yang, 2012). HQ exposure has been associated with oxidative damage and could affect the oxidative balance in other biological systems (Heluany et al., 2018b; Heluany et al., 2021; Peng et al. 2013; Pons et al. 2010). Indeed, here we confirmed the pro-oxidative effect of HQ on the articular chondrocytes, which could contribute to the overall phenotypic changes induced by the xenobiotic on these cells.

Interleukin 1β (IL-1β) is a major pro-inflammatory trigger in OA that stimulates catabolic changes, suppresses anabolic pathways and decreases matrix synthesis (Jenei-Lanzl, Meurer and Zaucke, 2019). In our study, IL-1β and HQ had a synergistic effect in reducing proteoglycan content and in promoting oxidative stress in the articular chondrocytes, suggesting that exposure to cigarette smoke and environmental pollutants could potentiate inflammatory processes involved in the degradation of the articular cartilage in OA. Interestingly, similar synergistic effects were observed in synoviocytes derived from RA-patients co-stimulated with HQ and TNF-α (Heluany et al., 2021).

AhR is a ligand-dependent transcription factor that translocates to the nucleus upon activation by xenobiotics and pollutants (Julliard, Fechner and Mezrich, 2014). Here we confirmed that after HQ exposure, AhR translocates from cytoplasm to the nucleus, forms a heterodimer with the AhR nuclear translocator (ARNT) and promotes the transcription of target genes such as Cyp1a1. The activation of the AhR pathway has been shown to mediate several detrimental effects, such as aggravation of articular diseases, cancer promotion and endocrine disruption (Denison and Nagy, 2003; Nakahama et al., 2011; Nguyen et al., 2015). Several studies showed that this receptor plays a major role in exacerbating rheumatoid arthritis in smokers (Kobayashi et al., 2008; Nguyen et al., 2015; Talbot et al., 2018). We have also previously showed the HQ-mediated cytotoxicity in joint diseases through the activation of the AhR pathway (Heluany et al., 2018a,b; Heluany et al., 2021). Here we show that AhR and its downstream effectors are overexpressed by HQ in articular chondrocytes, and mediate the HQ catabolic effects, suggesting that xenobiotics and pollutants can directly influence the health of the articular cartilage.

In summary, our data demonstrated a clear detrimental effect of HQ on articular cartilage homeostasis and shed novel insight on how environmental pollutants can exacerbate the degenerative effect of pro-inflammatory mechanisms underlying the onset of articular diseases such as OA.

## Conflict of interests

*The authors declare that the research was conducted in the absence of any commercial or financial relationships that could be construed as a potential conflict of interest*.

## Author Contributions

CSH: Conceptualization, data collection, analyses, interpretation and manuscript writing, revision and proof-read. ADP: Data collection, manuscript proof-read. ND: Data collection, manuscript proof-read. SHPF: Conceptualization, supervision, interpretation and manuscript writing revision and proof-read. GN: Conceptualization, supervision, interpretation and manuscript writing, revision and proof-read.

## Funding

This work was supported by the following grants: Fundação de Amparo à Pesquisa do Estado de São Paulo [FAPESP – C.S.H. – grant number 2019/24639-7], the Medical Research Council MR/S008608/1 and the Academy of Medical Sciences SBF004\1112.

## Notes

### Competing Interest Statement

The authors have declared no competing interest.

## References

Akkiraju, H., and Nohe, A. (2015). Role of Chondrocytes in Cartilage Formation, Progression of Osteoarthritis and Cartilage Regeneration. J. Dev. Biol. 3:177–192. doi:10.3390/jdb3040177.

Abramson, S.B. (2008). Osteoarthritis and nitric oxide. Osteoarthr. Cartil. 16:S15–S20. doi:10.1016/S1063-4584(08)60008-4.

Amin, A., and Abramson, S.B. (1998). The role of nitric oxide in articular cartilage breakdown in osteoarthritis. Curr. Opin. Rheumatol. 10:263–268. doi:10.1097/00002281-199805000-00018.

Amim, S., Niu, J., Guermazi, A., Grigoryan, M., Hunter, D.J., Clancy, M. et al. (2007). Cigarette smoking and the risk for cartilage loss and knee pain in men with knee osteoarthritis. Ann. Rheum. Dis. 66:18–22.

Arner, E.C. (2002). Aggrecanase-mediated cartilage degradation. Curr. Opin. Pharmacol. 2(3):322–329. doi:10.1016/s1471-4982(02)00148-0.

Berenbaum, F. (2013). Osteoarthritis as an inflammatory disease (osteoarthritis is not osteoarthrosis!). Osteoarthr. Cartil. 21:16–21. doi:10.1016/j.joca.2012.11.012.

Bodnar, J. A., Morgan, W.T., Murphy, P.A., and Ogden, M.W. (2012). Mainstream smoke chemistry analysis of samples from the 2009 US cigarette market. Regul. Toxicol. Pharmacol. 64(1):35–42. doi:10.1016/j.yrtph.2012.05.011.

Bortoluzzi, A. Furini, and F. Sciré, C.A. (2018). Osteoarthritis and its management - Epidemiology, nutritional aspects and environmental factors. Autoimmunity Rev. 17(11):07–1114. doi:10.1016/j.autrev.2018.06.002.

Cho, J. Y. (2008). Suppressive effect of hydroquinone, a benzene metabolite, on in vitro inflammatory responses mediated by macrophages, monocytes, and lymphocytes. Mediators Inflamm. 2008:298010. https://doi:10.1155/2008/298010.

Davies-Tuck, M.L., Wluka, A.E., Forbes, A., Wang, Y., English, D.R., Giles, G.G., and Cicuttini, F. (2009). Smoking is associated with increased cartilage loss and persistence of bone marrow lesions over 2 years in community-based individuals. Rheumatology 48:1227–1231. doi:10.1093/rheumatology/kep211.

De Bari, C., Dell’Accio, F., and Luyten, F.P. (2001). Human periosteum-derived cells maintain phenotypic stability and chondrogenic potential throughout expansion regardless of donor age. Arthritis Rheum. 44:88–95. doi:10.1002/1529-0131(200101)44:1<85::AID-ANR12>3.0.CO;2-6.

Dell’Accio, F., De Bari, C., and Luyten, F. P. (2001). Molecular markers predictive of the capacity of expanded human articular chondrocytes to form stable cartilage in vivo. Arthritis Rheum. 44(7):1608–19. doi:10.1002/1529-0131(200107)44:7<1608::AID.

Denison, M.S., and Nagy, S.R. (2003). Activation of the aryl hydrocarbon receptor by structurally diverse exogenous and endogeneous chemicals. Annu. Rev. Pharmacol. Toxicol. 43:309–34. doi:10.1146/annurev.pharmtox.43.100901.135828.

Farndale, R.W., Sayers, C.A., and Barrett, A. J. (1982). A direct spectrophotometric microassay for sulfated glycosaminoglycans in cartilage cultures. Connect Tissue Res. 9:247–248. doi:10.3109/03008208209160269.

Felson, D.T., and Zhang, Y. (2015). Smoking and osteoarthritis: a review of the evidence and its implications. Osteoarthr. Cartil. 23(3):331–3. doi:10.1016/j.joca.2014.11.022.

Fogelholm, R.R., and Alho, A.V. (2001). Smoking and intervertebral disc degeneration. Med. Hypotheses 56(4):537–9. doi:10.1054/mehy.2000.1253.

Goldberg, M.S., Scott, S.C., and Mayo, N. (2000). A review of the association between cigarette smoking and the development of nonspecific back pain and related outcomes. Spine 25(8):995–1014. doi:10.1097/00007632-200004150-00016.

Heluany, C.S., Kupa, L.V.K., Viana, M.N., Fernandes, C.M., and Farsky, S.H.P. (2018a). Hydroquinone exposure worsens the symptomatology of rheumatoid arthritis. Chem. Biol. Interact. 291:120–127. doi:10.1016/j.cbi.2018.06.016.

Heluany, C.S., Kupa, L.V.K., Viana, M.N., Fernandes, C.M., Silveira, E.L.V. and Farsky, S.H.P. (2018b). In vivo exposure to hydroquinone during the early phase of collagen-induced arthritis aggravates the disease. Toxicology 481:22–30. doi:10.1016/j.tox.2018.06.010

Heluany, C.S., Donate, P.B., Schneider, A.H., Fabris, A.L., Gomes, R. A., Villas-Boas, I., et al. (2021). Hydroquinone Exposure Worsens Rheumatoid Arthritis through the Activation of the Aryl Hydrocarbon Receptor and Interleukin-17 Pathways. Antioxidants 10:929. doi:10.3390/antiox10060929

Henrotin, Y.E., Bruckner, P., and Pujol, J.P. (2003). The role of reactive oxygen species in homeostasis and degradation of cartilage. Osteoarthr. Cartil. 11:747e55. doi:10.1016/s1063-4584(03)00150-x.

Jenei-Lanzl, Z., Meurer, A., and Zaucke, F. (2019). Interleukin-1β signalling in osteoarthritis – chondrocytes in focus. Cellular Sig. 53:212–223. doi:10.1016/j.cellsig.2018.10.005

Julliard, W., Fechner, J.H., and Mezrich, J.D. (2014). The aryl hydrocarbon receptor meets immunology: friend or foe? A little of both. Front. Immunol. 5:458. doi:10.3389/fimmu.2014.00458.

Karaliotas, G.I., Mavridis, K., Scorilas, A., and Babis, G.G. (2015). Quantitative analysis of the mRNA expression levels of BCL2 and BAX genes in human osteoarthritis and normal articular cartilage: An investigation into their differential expression. Mol. Med. Rep. 12:4514–4521. doi:10.3892/mmr.2015.3939

Kobayashi, S., Okamoto, H., Iwamoto, T., Toyama, T., Tomatsu, T., Yamanaka, H. et al. (2008). A role for the aryl hydrocarbon receptor and the dioxin TCDD in rheumatoid arthritis. Rheumatology 47:1317–1322. https://doi:10.1093/rheumatology/ken259

Lee, J. S., Yang, J. S., Yang, E. J., and Kim, I. S. (2012). Hydroquinone-induced apoptosis of human lymphocytes through caspase 9/3 pathway. Mol. Biol. Rep. 39(6):6737–43. doi:10.1007/s11033-012-1498-y

Lee, H.G., and Yang, J.H. (2012). PCB126 induces apoptosis of chondrocytes via ROS-dependent pathways. Osteoarthr. Cartil. 20(10):1179–85. doi:10.1016/j.joca.2012.06.004.

Li, J., Jiang, S., Chen, Y., Ma, R., Chen, J., Qian, S. et al. (2018). Benzene metabolite hydroquinone induces apoptosis of bone marrow mononuclear cells through inhibition of β-catenin signaling. Toxicol. In Vitro 46:361–369. doi:10.1016/j.tiv.2017.08.018

Li, J., Xing, X., Zhang, X., Liang, B., He, Z., Gao, C. et al. (2018). Enhanced H3K4me3 modifications are involved in the transactivation of DNA damage responsive genes in workers exposed to low-level benzene. Environ. Pollut. 234:127–135. https://doi:10.1016/j.envpol.2017.11.042

Ma, Q., and Baldwin, K. T. (2000). 2,3,7,8-tetrachorodibenzo-p-dioxin-induced degradation of aryl hydrocarbon receptor (AhR) by the ubiquitin-proteasome pathway. Role of the transcription activation and DNA binding. J. Biol. Chem. 275(12):8432–8. https://doi:10.1074/jbc.275.12.842.

Mao, J., Dai, W., Zhang, S., Sun, L., Wang, H., Gao, Y. et al. (2019). Quinone-thioether metabolites of hydroquinone play a dual role in promoting a vicious cycle of ROS generation: in vitro and in silico insights. Arch. Toxicol. 93(5):1297–1309. doi:10.1007/s00204-019-02443-4

McGregor, D. (2007). Hydroquinone: an evaluation of the human risks from its carcinogenic and mutagenic properties. Crit. Rev. Toxicol. 37:887–914. doi:10.1080/10408440701638970

Mobasheri, A., Rayman, M.P., Gualillo, O., Sellam, J., Van Der Kraan, P., and Fearon, U. (2017). The role of metabolism in the pathogenesis of osteoarthritis. Nat. Rev. Rheumatol. 13:302. doi:10.1038/nrrheum.2017.50.

Nakahama, T., Kimura, A., Nguyen, N.T., Chinen, I., Hanieh, H., Nohara, K. et al. (2011). Aryl hydrocarbon receptor deficiency in T cells suppresses the development of collagen-induced arthritis. Proc. Natl. Acad. Sci. USA 108:14222–14227. doi:10.1073/pnas.1111786108

Nalesso, G., Sherwood, J., Bertrand, J., Pap, T., Ramachandran, M., De Bari, C. et al. (2011). WNT-3A modulates articular chondrocyte phenotype by activating both canonical and noncanonical pathways. J. Cell Biol. 193:551–64. doi:10.1083/jcb.201011051.

Nalesso, G., Thomas, B.L., Sherwood, J.C., Yu, J., Addimanda, O., Eldrige, S.E. et al. (2017). WNT16 antagonises excessive canonical WNT activation and protects cartilage in osteoarthritis. Ann. Rheum. Dis. 76:218–226. doi:10.1136/annrheumdis-2015-208577.

Nguyen, N.T., Nakahama, T., Nguyen, C.H., Tran, T.T., Le, V.S., Chu, H.H., and Kishimoto, T. Aryl hydrocarbon receptor antagonism and its role in rheumatoid arthritis. J. Exp. Pharmacol. 7:29–35. doi:10.2147/JEP.S63549

Peng, C., Arthur, D., Liu, F., Lee, J., Xia, Q., Lavin, M.F., and Ng, J.C. (2013). Genotoxicity of hydroquinone in A549 cells. Cell Biol. Toxicol. 29(4):213–227. doi:10.1007/s10565-013-9247-0

Poole, A. R., Kobayashi, M., Yasuda, T. et al. (2002). Type II collagen degradation and its regulation in articular cartilage in osteoarthritis. Ann. Rheum. Dis. 61:ii78–ii81. doi:10.1136/ard.61.suppl_2.ii78.

Pons, M., Cousins, S.W., Csaky, K.G., Striker, G., and Marin-Castaño, M.E. (2010). Cigarette smoke-related hydroquinone induces filamentous actin reorganization and heat shock protein 27 phosphorylation through p38 and extracellular signal-regulated kinase 1/2 in retinal pigment epithelium: implication for age-related macular degeneration. Am. J. Pathol. 177(3):1198–213. doi:10.2353/ajpath.2010.091108

Racunica, T.L., Szramka, M., Wluka, A.E., Wang, Y., English, D.R., Giles, G.G. et al. (2007). A positive association of smoking and articular knee joint cartilage in healthy people. Osteoarthr. Cartil. 15:587–590. doi:10.1016/j.joca.2006.12.005.

Stabbert, R., Dempsey, R., Diekmann, J., Euchenhofer, C., Hagemeister, T., Haussmann, H-J. et al. (2017). Studies on the contributions of smoke constituents, individually and in mixtures, in a range of in vitro bioactivity assays. Toxicol. Vitr. 42:222–246. doi:10.1016/j.tiv.2017.04.003

Sellan, J., Berenbaum, F. (2010). The role of synovitis in pathophysiology and clinical symptoms of osteoarthritis. Nat. Rev. Rheumatol. 6(11):625–35. doi:10.1038/nrrheum.2010.159.

Stampfli, M.R., and Anderson, G.P. (2009). How cigarette smoke skews immune responses to promote infection, lung disease and cancer. Nat. Rev. Immunol. 9:377–384. doi:10.1038/nri2530

Suantawee, T., Tantavisut, S., Adisakwattana, S., Tanavalee, A., Yuktanandana, P., Anomasiri, W. et al. (2013). Oxidative stress, vitamin e, and antioxidant capacity in knee osteoarthritis. J. Clin. Diagn. Res. 7:1855–1859. doi:10.7860/JCDR/2013/5802.3333.

Talbot, J., Peres, R.S., Pinto, L.G., Oliveira, R.D.R., Lima, K.A., Donate, P.B. et al. (2018). Smoking-induced aggravation of experimental arthritis is dependent of aryl hydrocarbon receptor activation in Th17 cells. Arthritis Res. Ther. 20:119. doi:10.1186/s13075-018-1609-9

Thomas, C.M., Fuller, C.J., Whittles, C.E., and Sharif, M. (2007). Chondrocyte death by apoptosis is associated with cartilage matrix degradation. Osteoarthr. Cartil. 15:27–34. doi:10.1016/j.joca.2006.06.012.

